# Using targeted re-sequencing for identification of candidate genes and SNPs for a QTL affecting the pH value of chicken meat

**DOI:** 10.1101/017186

**Authors:** Xidan Li, Xiaodong Liu, Javad Nadaf, Elisabeth Le Bihan-Duval, Cécile Berri, Ian C. Dunn, Richard Talbot, Dirk-Jan de Koning

**Affiliations:** Swedish University of Agricultural Sciences, SE-750 07 Uppsala, Sweden; The Linnaeus Center for Bioinformatics, Uppsala University, SE-75124 Uppsala, Sweden; INRA, UR83, Recherches Avicoles, Nouzilly, 37380, France; Roslin Institute and R(D)SVS, University of Edinburgh, Midlothian, EH25 9RG, United Kingdom

**Author notes:** To whom correspondence should be addressed: Prof. Dirk-Jan de Koning Department of Animal Breeding and Genetics Swedish University of Agricultural Sciences Box 7023, SE-750 07 Uppsala, Sweden.

**Keywords:** Chicken muscle, pH values, QTL, Next Generation Sequencing, SNPs, non-synonymous SNPs

## Abstract

Using targeted genetical genomics, a QTL affecting the initial post-mortem pH value of chicken breast muscle (*Pectoralis major*) on chromosome 1 (GGA1) was recently fine-mapped. Thirteen genes were present in the QTL region of about 1 Mb.

In this study, ten birds that were inferred to be homozygous for either the high (QQ) or low (qq) QTL allele were selected for re-sequencing. After enrichment for 1 Mb around the QTL region, > 200 x coverage for the QTL region in each of the ten birds was obtained. We used custom tools to identify putative causal mutations in the mapped QTL region from next generation sequence data. Four non-synonymous SNPs differentiating the two QTL genotype groups were identified within four local genes (*PRDX4, EIF2S3, PCYT1B and E1BTD2*). These were defined to be most likely candidate SNPs to explain the QTL effect. Moreover, 29 consensus SNPs were detected within gene-related regions (UTR regions and splicing sites) for the QQ birds and 26 for the qq birds. These could also play a role explaining the observed QTL effect.

The results provide an important step for prioritizing among a large amount of candidate mutations and significantly contribute to the understanding of the genetic mechanisms affecting the initial post-mortem pH value of chicken muscle.

## Introduction

Meat quality traits such as color, water holding capacity and texture are important in poultry and are determined by both genetic and environmental factors. The post-mortem pH drop is a key factor affecting these meat quality traits (Berri *et al.* 2007). Meat with a low pH value tends to be pale in color with low water holding capacity, which has a critical impact on quality of further processed products (Dransfield 1999; Alnahhas *et al.*, 2014). Previous studies have shown that ultimate pH of meat is highly dependent upon the amount of glycogen in the muscle (Le Bihan-Duval *et al.* 2008). This glycogen is rapidly depleted in the muscle while the birds are exposed to stress (Ngoka and Froning, 1982) and exhibit high physical activity (Debut *et al.*, 2005) prior to slaughter. Therefore, pre-slaughter stress may be associated with variation in the initial rate of drop in pH (Chabault *et al.*, 2012). At the genetic level, a recent fine mapping study has identified a QTL region affecting muscle pH measured 15 minutes post-mortem (pH15) using an F_2_ intercross between High Growth (HG) and Low Growth (LG) chicken lines (Nadaf *et al.* 2007, 2014). The HG and LG lines are being bi-directionally selected on body weight at nine week of age, where the HG lines show lower pH15 values and the LG lines show higher pH15 values. Identifying the causal genetic component underlying this QTL region would advance our understanding of genetic architecture of poultry muscle metabolism.

In this study, ten birds that were homozygous at the targeted QTL (QQ vs. qq) were selected for re-sequencing, where QQ corresponds to alleles from the HG lines and qq corresponds to alleles from the LG lines (Nadaf *et al.* 2014). Based on the bioinformatics analysis of the mapped QTL region and SNPs detected by Next Generation Sequencing (NGS) data, we report a number of candidate genes and SNPs.

## Materials and methods

### Study samples

In this study, we selected 10 birds from the F2 with inferred QTL genotypes of QQ (5 birds, homozygous for HG line alleles) or qq (5 birds, homozygous for LG line alleles). The selected individuals are a subset of the birds that were used for the targeted genetical genomics study by Nadaf *et al.* (2014).

## Re-mapping the QTL region in the galGal 4.0 genome

The previously fine-mapped QTL region, which spanned less than 1Mbps on chromosome 1, was re-sequenced based on the reference genome galGal 3.0. However, with the recently assembled reference genome galGal 4.0, the updated gene information had some differences compared to galGal 3.0 for our target region. Therefore, we re-mapped the QTL region onto the genome assembly galGal 4.0, where sequence of the fine-mapped QTL region on genome assembly galGal 3.0 was blasted against the genome sequence galGal 4.0 (Zhang *et al.* 2000). As a result, the new QTL region of around 1.2 Mbps has been mapped to chr1: 117000000 - 118200000 (on galGal4.0, see Figure 1).

**Figure 1.**
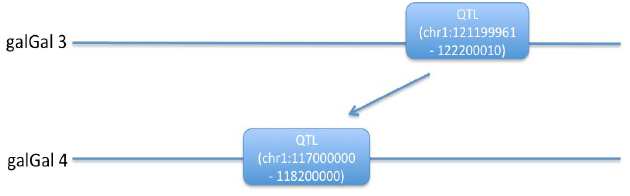
Re-mapping the QTL region from the galGal 3.0 genome to the galGal 4.0 genome.

## Next Generation Sequencing analysis

Target enrichment analyses and next generation sequencing were performed by Edinburgh Genomics (https://genomics.ed.ac.uk/). The previously mapped QTL regions were re-sequenced based on the reference genome galGal 3.0. In order to get high coverage in the QTL region, Agilent SureSelect (Agilent, Santa Clara, USA) target enrichment system was used to perform DNA sequence enrichment. SureSelect technology is designed to isolate a subset (up to 24Mb) of a genome region or regions for high throughput sequencing. Sequencing was done using Illumina paired-end libraries with about 150 bps for each pair of read and an average insert size of 300 bps.

The Illumina Genome Analyzer Analysis Pipeline was used to produce paired sequence files containing reads for each sample in Illumina FASTQ format. All sequences have been deposited in the Sequence Read Archive (SRA, http://www.ncbi.nlm.nih.gov/sra) under accession number SRP051545. The sequences alignments were done using bowtie (Langmead *et al.* 2009) based on the re-mapped QTL region on the genome galGal 4.0, where about 500X coverage for each of ten homozygote genotypes was achieved. Further, samtools (Li *et al.* 2009) was used to perform SNP-calling, and the resulting VCF files were used for further candidate gene analysis (workflow provided in Supplementary File 1).

### Detecting potential causative genes and SNPs corresponding to the genotypic differences between two genotype groups

The QTL region affecting the pH15 value of chicken meat has been previously fine mapped (Nadaf *et al.* 2014). Although QTL analysis has been shown to be useful to detect the genetic variations related to the common complex trait and genotype, the ability to detect the causative functional genes and SNPs is scarce. In this study, we develop a new strategy to identify the functional SNPs in known genes in a defined QTL region, which could prioritize the most promising SNPs among a large pool of candidates (workflow and subroutines presented in Supplementary File2). From the VCF files, the gene information such as gene’s transcript id, the coordinate of RNA splicing sites, and UTR regions in the re-mapped QTL region were retrieved from Ensembl (Flicek *et al*. 2013). Next, SNPs located within the non-coding regions such as RNA splicing sites and UTR regions were identified and recorded as potential factors affecting gene expression. SNPs in the coding regions that were non-synonymous mutations were evaluated and ranked according to their effect on the protein function by PASE (Li *et al*. 2013) (Figure 2). Finally, the identified SNPs from each sample were categorized according to their genotypes. The SNPs that are present in, almost, every sample of one QTL genotype group (e.g. QQ) and absent in the other QTL genotype group are called ‘consensus SNPs. These consensus SNPs were chosen as candidate SNPs and the corresponding genes were prioritized for future analyses.

**Figure 2.**
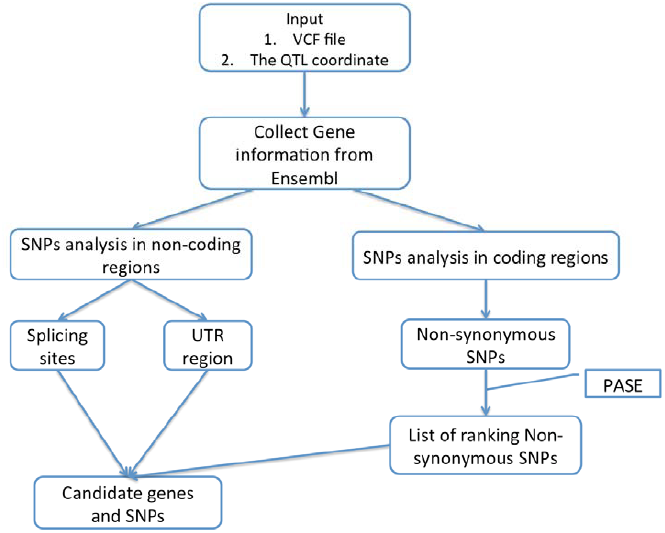
Schematic overview of the workflow to identify the candidate genes and SNPs

## Results and Discussion

In this study we identified 60 gene related SNPs that separated the two QTL genotype groups (Table 1). Among these, four non-synonymous SNPs (nsSNPs) were identified within four genes (Table 2). One consensus nsSNP located in the gene PRDX4 was detected in four out of five QQ birds. Previous studies show that the pH15 value of chicken meat is associated with straightening up and wing flapping at the slaughter shackle line (Berri, *et al*. 2005 and Nadaf *et al.* 2007). These activities at the slaughter line could directly result in fatigue of the muscle by the production of oxidants. Therefore, oxidative stress has been suggested to contribute to the pH15 value of chicken meat (Nadaf *et al*. 2014). Interestingly, PRDX4, as one member of the antioxidant enzyme family, plays a crucial role in the process of protecting muscle cells against oxidative stress (Novelli *et al. 1991*; Leyens *et al.* 2003). Our previous study has shown that the expression level of PRDX4 is significantly different between the QQ and qq genotype groups (Nadaf *et al*. 2014). Therefore, PRDX4 remains a strong candidate gene for this QTL. The three other nsSNPs were also detected within three genes: *EIF2S3, PCYT1B* and *E1BTD2* (Table 2). *EIF2S3* has been described to contribute to the early stage of protein synthesis by interacting with GTP and initiator tRNA to form a ternary complex and binding to a 40S ribosomal subunit (Entrez Gene: EIF2S3). *PCYT1B* is an enzyme, and involved in the regulation of phosphatidylcholine biosynthesis. Several alternatively spliced transcript variants encoding different isoforms have been found for this gene (Wang *et al.* 2005). Another gene, *E1BTD2* is uncharacterized. By using BLAST, the predicted function of *E1BTD2* was “organic solute transporter subunit alpha-like”, which is involved in the activity of transportation from the endoplasmic reticulum to the plasma membrane (Dawson *et al.* 2005). In summary, these genes are all involved in energy metabolism and could potentially explain the observed QTL effects.

**Table 1.**
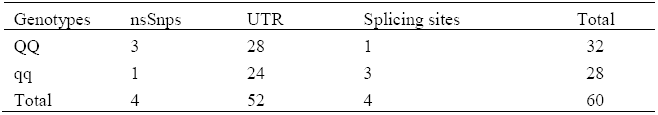
Number of consensus SNPs in UTR and Splicing site differentiating the QQ and qq genotypes for a QTL affecting pH after slaughter on chicken chromosome 1. Details of all SNP locations are in Supplementary Table 1.

**Table 2.** Consensus non-synonymous SNPs differentiating the QQ and qq genotypes for a QTL affecting pH after slaughter on chicken chromosome 1.

| Gene ID | Genotype | Non-synonymous SNPs |  |  |
| --- | --- | --- | --- | --- |
|  |  | Coordinate | Codon (Ref/Alt) | Amino Acid Substitution (Ref/Alt) |
| PRDX4 | QQ | 117796698 | ACT/CCT (4/5) <sup>1</sup> | T/P |
| EIF2S3 | QQ | 117669996 | GGT/AGT | G/S |
| E1BTD2 | QQ | 117608741 | TTT/ATT | F/I |
| PCYT1B | qq | 117503317 | GCA/GGA | A/G |

<sup>1</sup> *The consensus nsSNP located in the gene PRDX4 has been detected in four out of five QQ*
*genotype samples*

Moreover, 29 consensus SNPs were detected within gene-related regions (UTR regions and splicing sites) for the QQ birds and 26 for the qq birds (Table 1). Because variations in regulatory elements are expected to contribute to variation in complex traits, these consensus SNPs could affect the expression of target genes and contribute to the observed QTL effect. For example, one consensus SNP, present in the QQ group, was identified at a canonical splicing site of *ACOT9*, which could result in inhibition of splicing and have a decisive impact on its function. In addition, *ACOT9* was differentially expressed in both the microarray and Real-time PCR analysis between QQ and qq groups in the previous study (Nadaf *et al.* 2014) (supplementary Table 1). Thus, *ACOT9* is another strong candidate gene for the QTL effect.

In the current study we used target enrichment to get high coverage of our QTL region in the sequence data. This has been successful in terms of creating high average coverage but there are also some drawbacks: Because the target enrichment was based on 1 Mb on genome build galGal 3.0, the re-mapped region of 1.2 Mb on genome build galGal 4.0 means that we cannot have continuous sequence coverage of the QTL region. Furthermore, the target enrichment process ignores repetitive DNA sequences in the target region causing addition problems for having continuous coverage. These factors mean that the current re-sequencing may have missed polymorphisms in the target region. Given the current prices for sequencing, we would recommend to do a full genome sequence of these target birds rather than only the QTL region.

In conclusion, the results confirm the previous study conclusion that suggests *PRDX4* and *ACOT9* are strong candidates for the QTL affecting the pH15 value of chicken meat. The list of plausible candidate genes and mutations from the present study will facilitate further verification and experimental evaluation. With the current study we have fully exploited the potential for fine mapping in the divergent F2 cross. To differentiate among the current list of candidate genes and SNPs we will need additional experiments. For instance, the candidate SNPs could be tested on a broader panel of birds from different breeds to evaluate their putative effect. Thus, an improved understanding of the genetic basis of variation in pH values will assist selection of breeding birds.

## Acknowledgments

JN was funded by the Marie Curie Host Fellowships for Early Stage Research Training, as part of the 6th Framework Programme of the European Commission. DJK, RT and ID acknowledge support from the BBSRC through The Roslin Institute ISPG. Edinburgh Genomics, at The University of Edinburgh is partly supported through core grants from NERC (R8/H10/56), MRC (MR/K001744/1) and BBSRC (BB/J004243/1). The sequencing results are part of the SABRE research project that has been co-financed by the European Commission, within the 6th Framework Programme, contract No. FOOD-CT-2006-016250. This publication represents the views of the authors, not the European Commission.

